# Palmitate-induced toxicity is associated with impaired mitochondrial respiration and accelerated oxidative stress in cultured cardiomyocytes: the critical role of Coenzyme Q_9/10_

**DOI:** 10.1101/830331

**Authors:** Phiwayinkosi V. Dludla, Sonia Silvestri, Patrick Orlando, Sithandiwe E. Mazibuko-Mbeje, Rabia Johnson, Fabio Marcheggiani, Ilenia Cirilli, Christo J.F. Muller, Johan Louw, Nireshni Chellan, Nnini Obonye, Bongani B. Nkambule, Luca Tiano

**Author notes:** **Corresponding author:** Phiwayinkosi V. Dludla, PhD, Biomedical Research and Innovation Platform, South African Medical Research Council, Tygerberg 7505, South Africa.

## Abstract

Impaired mitochondrial function concomitant to enhanced oxidative stress-induced damage are well established mechanisms involved in hyperlipidemia-induced cardiotoxicity. Coenzyme Q9/10 (CoQ) is known to be a critical component of the mitochondrial electron transport chain that efficiently supports the process of bioenergetics in addition to its antioxidant activities. However, there is very limited information on the direct effect of myocardial lipid overload on endogenous CoQ levels in association with mitochondrial respiration and oxidative stress status. Here, such effects were explored by exposing H9c2 cardiomyocytes to various doses (0.15 to 1 mM) of palmitate for 24 hours. The results demonstrated that palmitate doses ≥ 0.25 mM are enough to impair mitochondrial respiration and cause oxidative stress. Although endogenous CoQ levels are enhanced by palmitate doses ≤ 5 mM, this is not enough to counteract oxidative stress, but is sufficient to maintain cell viability of cardiomyocytes, suggesting a compensation mechanism. Palmitate doses > 5 mM caused severe mitochondrial toxicity, including reduction of cell viability. Interestingly, enhancement of CoQ levels with the lowest dose of palmitate (0.15 mM) was accompanied by a significantly reduction of CoQ oxidation status, as well as low cytosolic production of reactive oxygen species. From the overall findings, it appears that CoQ response may be crucial to improve mitochondrial function and thus protect against hyperlipidemia-induced insult. These results further suggest that therapeutic agents that can stimulate endogenous levels of CoQ may be beneficial in protecting the myocardium against diabetes associated complications.

## 1. Introduction

Enhanced myocardial lipid accumulation is a crucial feature that greatly impact cardiac efficiency by promoting structural modifications that supersede cellular injury [1, 2]. In this case, a targeted approach to critically understand mechanisms implicated in lipid-induced cardiotoxicity would seem to be an ideal strategy to protect against diabetic cardiomyopathy (DCM). The latter describes a condition that specifically affects the myocardium in the absence of atherosclerosis, and has been another growing concern due to its long latent phase, during which the disease develops but is completely asymptomatic [1-3]. Certainly, enhanced cardiomyocyte triglyceride stores are observed in animal models of diabetes and obesity [4-6]. Also in hearts of patients with metabolic syndrome [7], enhanced intramyocyte lipid accumulation has been correlated with upregulated expression of adipogenic genes and reduced left ventricular ejection fraction, a well-known hallmark of DCM. Although the myocardium heavily relies on FFAs for energy generation, their exacerbated uptake and utilization can impair the efficiency of the electron transport chain and oxidative phosphorylation, in the process cause excess production of reactive oxygen species (ROS) [2, 8, 9]. Beyond impairing mitochondrial energetics [2, 9, 10], lipid-overload can severely deplete ubiquinone levels and thus contribute to oxidative stress-induced myocardial damage [11-13].

Ubiquinone, or rather Coenzyme Q_9/10_ (CoQ), is a naturally occurring fat-soluble component of the electron transport chain that participates in aerobic cellular respiration. In addition to its role as an electron transfer intermediate, CoQ also acts as a strong antioxidant in its reduced form, largely participates in the blockade of lipid peroxidation, thereby attenuate the reaction of hydroxyl (•OH) and superoxide (O_2_^• −^) radicals with lipid and protein molecules [13, 14]. The chain-breaking antioxidant properties of CoQ are important for blocking the initiation and propagation of oxidative stress [14]. Consequently, it is understood that conditions like diabetes can severely deplete endogenous CoQ levels across different tissues, including the myocardium, and thus accelerate oxidative stress-induced damage [15, 16]. While such information is accepted, data on the direct effect of lipid overload, as an *in vitro* model of DCM, on endogenous CoQ levels is still scanty. Thus, the current study explored a dose-dependent effect of palmitate, a known saturated FFA, on endogenous levels and oxidation status of CoQ in cultured H9c2 cardiomyocytes. Such data were compared to the impact of palmitate on mitochondrial respiration and cell viability status of these cardiomyocytes.

## 2. Materials and methods

### 2.1. Chemicals and reagents

Embryonic ventricular rat heart derived H9c2 cardiomyoblasts (CRL-1446) were from American Type Culture Collection (Manassas, USA); Dulbecco’s Modified Eagle’s Medium (DMEM), Penicillin-Streptomycin, Phosphate Buffered Saline (DPBS), and fetal bovine serum (FBS) were from Carlo Erba (Milano, Italy); Bradford and RC DC protein assay kit was from Bio-Rad Laboratories (Hercules, USA); 2’, 7’-dichlorfluorescein diacetate (DCFH-DA) and MitoSOX fluorescent dyes were from ThermoFisher Scientific (Waltham, USA); ViaCount was from Merck-Millipore (Darmstadt, Germany). Seahorse XF96 microplate plates, Seahorse XF Assay media, Seahorse XF base media without phenol red and Mito Stress Kits were from Agilent (Santa Clare, USA). All other chemicals including palmitate and 3-(4,5-Dimethylthiazol-2-yl)-2,5-Diphenyltetrazolium Bromide) (MTT) were obtained from Sigma-Aldrich (St. Louis, USA).

### 2.2. Cell culture and treatment conditions

H9c2 cardiomyoblasts were cultured in growth media containing DMEM supplemented with 10% FBS, 100 μg/mL penicillin and 100 μg/mL streptomycin overnight under standard tissue culture conditions (37 °C in humidified air and 5% CO_2_). Cells cultured at a seeding density of 25 000 cells/well in a 24 multi-well, and were maintained in growth media until confluency, whereas refreshed every second day. Confluent H9c2 cardiomyocyte were exposed to different palmitic acid doses (0.125, 0.25, 0.5, 0.75 and 1 mM) dissolved in 1% bovine serum albumin for 24 hours. Whereas, cardiomyocytes exposed to media containing 1% bovine serum albumin without any palmitate served as experimental controls.

### 2.3. CoQ quantification and oxidation status

The intracellular levels of CoQ_9_, which is the predominant form of ubiquinone in murine models [17], as well as the CoQ_10_, which is the well-known and predominant form in humans were quantified from treated H9c2 cardiomyocytes based on a previously described method [18]. Briefly, 50 μl cell suspension, approximately 200,000 cells, were extracted adding 250 μl isopropanol and vigorously mixing before centrifugation at 20,900 g for 2 minutes at 4°C. Thereafter, extracts were injected into a high-performance liquid chromatograph (HPLC) with electro-chemical detector (ECD) (Shiseido Co. Ltd.; Tokyo, Japan). This system is characterized by a post-separation reducing column (Shiseido CQR) that is capable of fully reducing eluted ubiquinone. CoQ_9/10_ oxidative status was expressed as percentage of ubiquinone/total CoQ_9/10_, while total CoQ_9/10_ content in H9c2 cardiomyocytes was expressed as μg/mg CoQ_9_ protein.

### 2.4. Mitochondrial and cytosolic ROS production assessment

Mitochondrial and cytosolic ROS production was evaluated using flow-cytometry by means of the reduced nerstian probe MitoSOX and DCFH-DA fluorescent dye, according to already described methods [19-21]. Briefly, treated cardiomyocytes or untreated control cells were incubated for 15 min or 30 min in a solution 5 μM of MitoSOX or DCFH-DA dye at 37°C. After trypsinization and washed cells were analyzed on a Guava Easycite flow cytometer (Millipore; Darmstadt, Germany).

### 2.5. Assessment of mitochondrial bioenergetics

Oxygen consumption rate (OCR) and extracellular acidification rates (ECAR) were measured using the XF96 Extracellular Flux analyzer (Agilent Technologies; Santa Clare, USA), as previously described [22]. OCR (pmol/min) was normalized with protein content, whereas OCR and ECAR were reported as absolute rates (pmoles/min for OCR and mpH/min for ECAR).

### 2.6. Cell viability and metabolic activity assays

Viability was estimated flow-cytometry using ViaCount Millipore cytotoxicity test as previously reported [19]. Briefly, the assay makes use of a mixture of cell membrane permeable and impermeable DNA-binding fluorescent probes, discriminated by red and yellow color, diluted 1:10 in PBS and used to stain approximately 20000 cells immediately before reading. The analysis of the distribution allows the discrimination of the percentage of cell debris (R-/Y-), live cells (R+/Y-) and dead cells (R+/Y+). On the other hand, a previously described method [23] making use of colorimetric measurement of the reduction of tetrazolium dye (MTT) to its insoluble formazan is a widely-used to measure metabolic activity of viable cells. Thereafter, absorbance was read at 570 nm using a BioTek ELx800 plate reader (Winooski, USA) and Gen 5 software for data acquisition.

### 2.7. Statistical analysis

Results were expressed as the mean of at least two independent biological experiments with each experiment containing at least three technical replicates. GraphPad Prism software version 6.0 (GraphPad Software, Inc., La Jolla, USA) was used to analyze data. Comparisons between groups were performed using one-way multivariate ANOVA followed by a Tukey post hoc test and Student *t*-test where appropriate. A p-value of < 0.05 was considered statistically significant.

## 3. Results

### 3.1. The dose-dependent effect of palmitate on endogenous CoQ levels

Due to mitochondrial abundance, cardiomyocytes are known to be generally rich in CoQ content, however CoQ_9_ levels are said to be more pronounced than that of CoQ_10_ as the current model is murine-derived cell line. Indeed, data presented in the current study showed that the levels of CoQ_9_ were much higher than that of CoQ_10_ (Fig. 1A-D). However, the trend of CoQ content was the same, showing a dose-dependent increase from 0.125 to 0.5 mM (p < 0.05 to p ≤ 0.001), then decreased with higher doses > 5 mM (p ≤ 0.05) of palmitate exposure, when compared to the experimental control (Fig. 1 A and B). Furthermore, although CoQ_10_ did not show any significance due to its relatively low content in the current experimental model, a similar trend was seen where a low dose palmitate (0.125 mM) demonstrated markedly reduced CoQ oxidation status when compared to the experimental control, while the higher doses, especially those ≥ 5 mM displayed complete oxidation status (Fig. 1 C and D).

**Figure 1.**
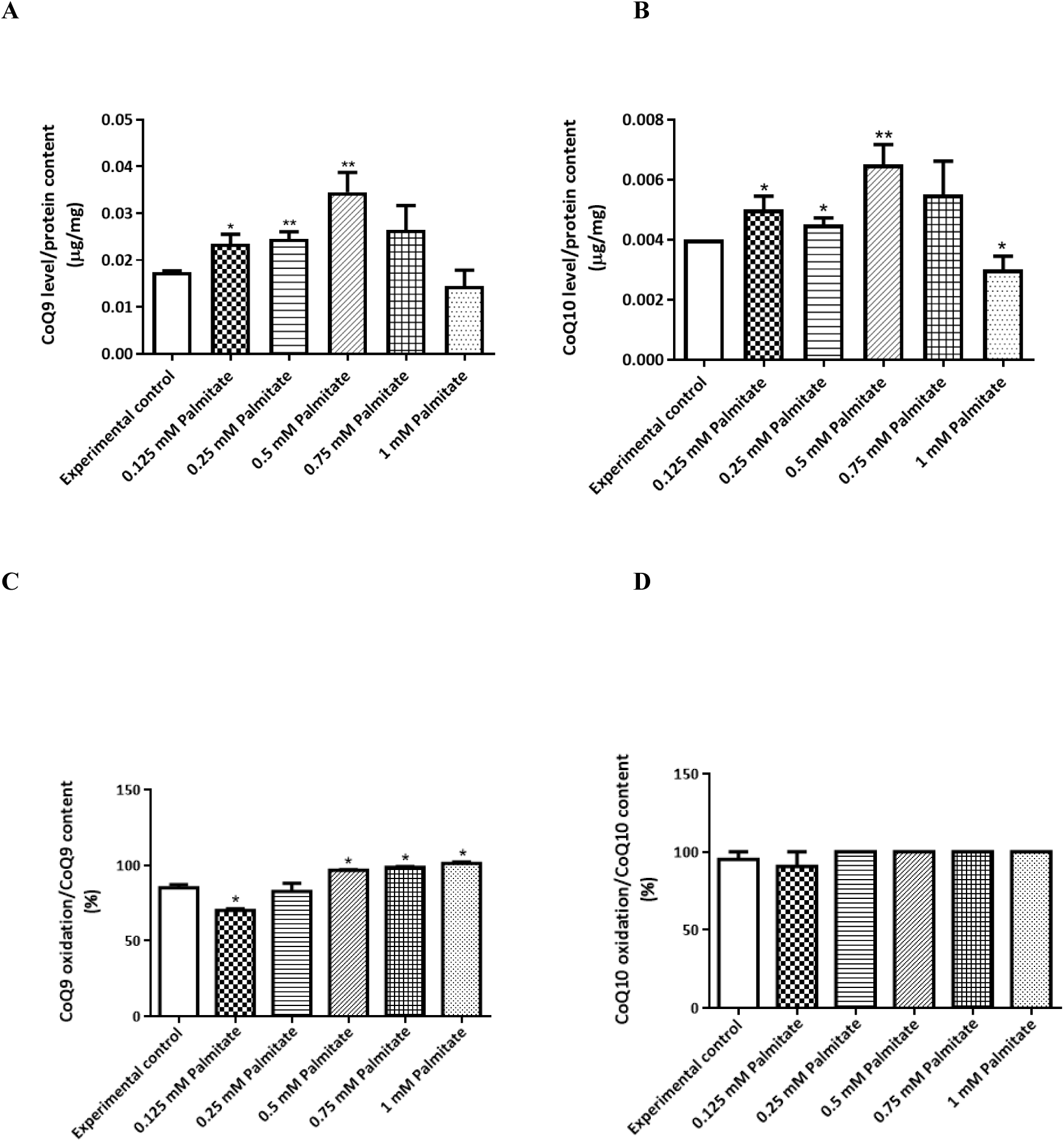
The dose-dependent effect of palmitate on endogenous Coenzyme Q_9/10_ (CoQ_9/10_) levels on cultured H9c2 cardiomyocytes. Briefly, H9c2 cardiomyocytes were exposed to various palmitate doses (0.125, 0.25, .0.75 and 1 mM) for 24 h before quantification of CoQ_9_ **(A)** and CoQ_10_ **(B)** levels, as well as CoQ_9_ **(C)** and CoQ_10_ **(D)** oxidation status. Results are expressed as the mean of at least two independent experiments relative to the experimental control. *p < 0.05, **p ≤ 0.01 versus experimental control.

### 3.2. The dose-dependent effect of palmitate on mitochondrial energetics

Efficiently controlled myocardial mitochondrial bioenergetics, especially the basal and maximal rate of respiration, is necessary for efficient production of energy in the form of ATP. Although did not significantly affect the basal oxygen consumption and maximal respiration rates, exposure of H9c2 cardiomyocytes to palmitate doses of 0.25 mM significantly reduced ATP production (Fig. 2A-D). Even worse, palmitate doses > 0.25 mM markedly (p ≤ 0.001) suppressed all components of mitochondrial bioenergetics, including ATP production, as well as oxygen consumption and maximal respiration rates (Fig. 2A-D).

**Figure 2.**
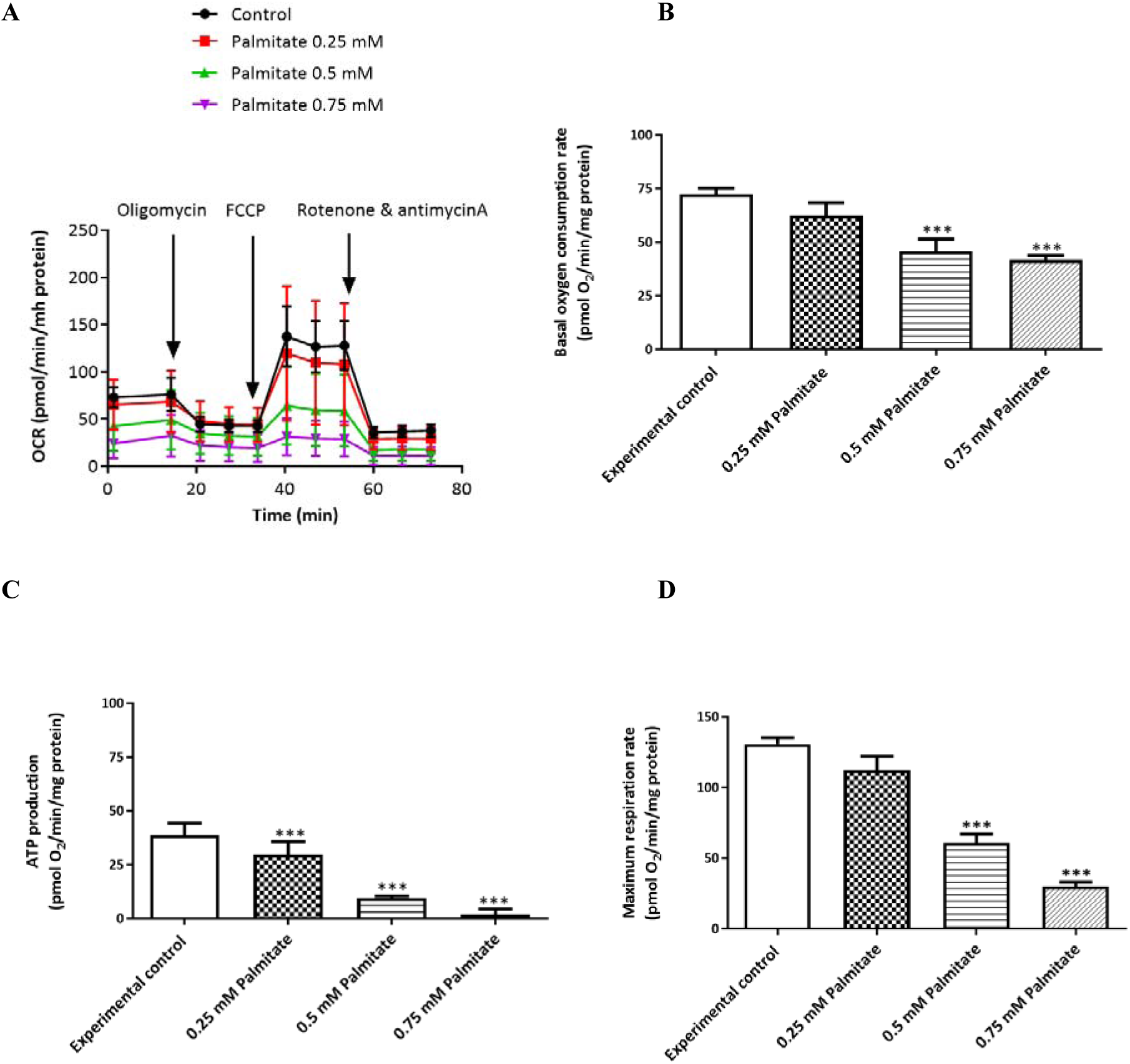
The dose-dependent effect of palmitate on mitochondrial energetics on cultured H9c2 cardiomyocytes. Briefly, H9c2 cardiomyocytes were exposed to various palmitate doses (0.125, 0.25, .0.75 and 1 mM) for 24 h before assessment of mitochondrial energetics. The graph shows oxygen consumption rate of all treatments (OCR) **(A)**, basal OCR **(B)**, ATP production **(C)**, and maximal respiration rate **(D)**. Results are expressed as the mean of at least two independent experiments relative to the experimental control. ***p ≤ 0.001 versus experimental control.

### 3.3. The dose-dependent effect of palmitate on mitochondrial and cytosolic ROS production

Both mitochondrial and cytosolic ROS production significantly contribute to oxidative stress-induced myocardial damage. Interestingly, there was no effect on mitochondrial ROS production observed with exposure of cardiomyocytes to the lowest dose of palmitate (0.125 mM) (Fig. 3 B). A slightly enhanced effect of ROS production, possible adaptive response, was observed with exposure of cardiomyocytes to palmitate doses between 0.25 and 0.5 mM. However, the current study also showed that doses > 5 mM of palmitate were enough to provoke enhanced mitochondrial and cytosolic production of ROS, when compared to the experimental control (p > 0.001) (Fig. 3 A and B).

**Figure 3.**
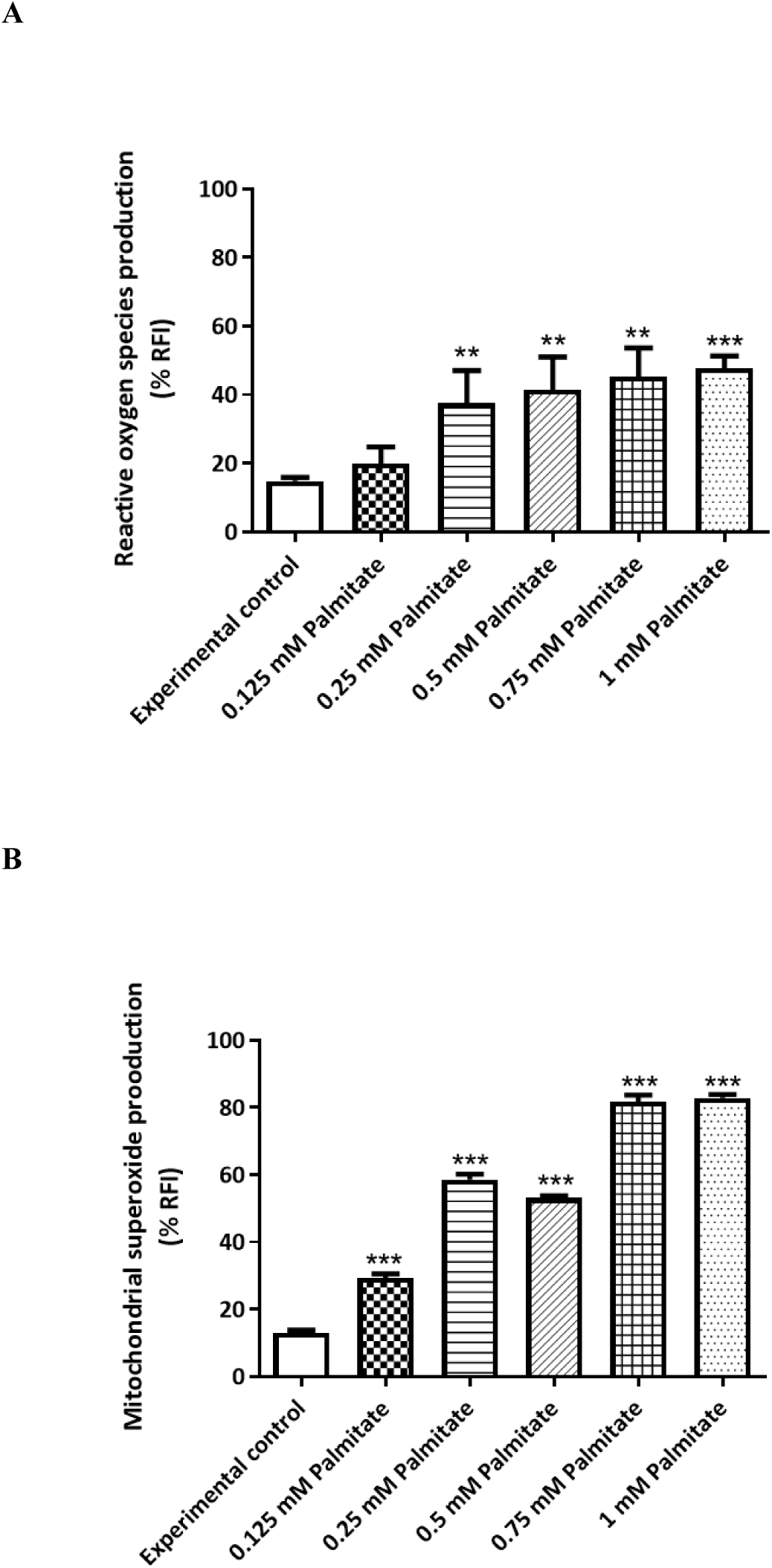
The dose-dependent effect of palmitate on mitochondrial and cytosolic production of reactive oxygen species (ROS). Briefly, H9c2 cardiomyocytes were exposed to various palmitate doses (0.125, 0.25, .0.75 and 1 mM) for 24 h before assessment of ROS production. The graphs represent cytosolic **(A)** and mitochondrial **(B)** ROS production. Results are expressed as the mean of at least two independent experiments relative to the experimental control. **p ≤ 0.01, ***p ≤ 0.001 versus experimental control.

### 3.4. The dose-dependent effect of palmitate on cell viability and metabolic activity

Accelerated cardiomyocyte damage because of lipid overload is a well-recognized phenomenon in a diabetic state. Here, data shows that an exposure of H9c2 cardiomyocytes to increasing doses of palmitate as a model of hyperlipidemia induced significantly reduced cell viability for concentrations of 0.25 mM (p < 0.05), and even worse for those > 0.25 mM (p ≤ 0.01). Such effects were even worse when metabolic activity was assessed, as for doses of 0.25 mM (p < 0.05) demonstrated significant reduction in metabolic activity, while even worse for doses ≥ 5 mM (p ≤ 0.001), when compared to the experimental control (Fig. 4 A and B).

**Figure 4.**
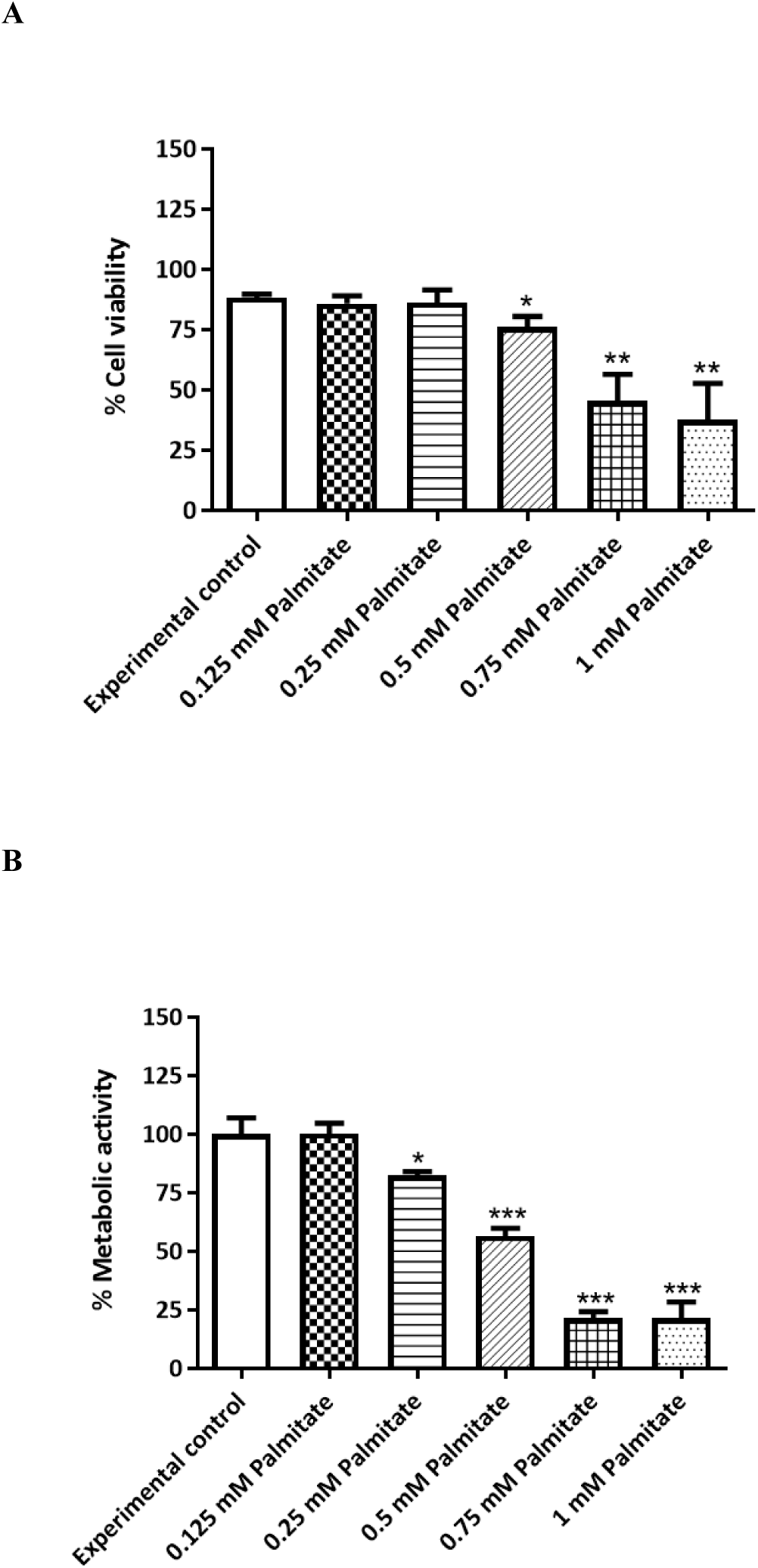
The dose-dependent effect of palmitate on cell viability and metabolic activity. Briefly, H9c2 cardiomyocytes were exposed to various palmitate doses (0.125, 0.25, .0.75 and 1 mM) for 24 h before assessment of cell viability and metabolic activity. The graphs represent cell viability **(A)** and metabolic activity **(B)**. Results are expressed as the mean of at least two independent experiments relative to the experimental control. *p < 0.05, **p ≤ 0.01, ***p ≤ 0.001 versus experimental control.

## 4. Discussion

To maintain its high energy demands for contractile function, the myocardium requires a continuous supply of substrates that are oxidized to produce ATP. In the healthy heart, this process is efficiently carried out by predominant oxidation of FFAs in the mitochondria. However, in conditions of stress, this process could be detrimental by accelerating mitochondrial ROS production and oxidative stress that normally leads to myocardial damage [2, 10]. In fact, lipid overload-induced mitochondrial ROS generation is reported in various models of metabolic syndrome and is concomitant to myocardial dysfunction [5, 7, 24]. Alternatively, in experimental models of hyperlipidemia, supplementation with CoQ has been shown to suppress oxidative stress by improving mitochondrial function [12, 25]. Beyond participation in oxidative phosphorylation, CoQ is known to be active scavenger of free radical species [14, 21], suggesting it could be used as a therapeutic to protect against DCM. Nevertheless, limited information is available on the direct effect of lipid overload on endogenous CoQ levels in association with mitochondrial function in the myocardium. Hence, the current study reports on a dose-dependent impact of lipid overload on mitochondrial energetics, endogenous CoQ levels and oxidative status in palmitate exposed H9c2 cardiomyoblasts.

The current study showed that exposure of H9c2 cardiomyocytes to compounding doses of palmitate could impair mitochondrial energetics as evident by reduced ATP production as a result of suppressed rates of oxygen consumption and maximal respiration. These results are of interest since it has already been established that myofibers obtained from diabetic patients also display reduced fatty acid-supported respiration, in addition to an increased content of myocardial triglycerides and increased mitochondrial H_2_O_2_ emission during oxidation of carbohydrate- and lipid-based substrates [26]. Also in our experimental model, the impairment of mitochondrial energetics was followed by accelerated mitochondrial and cytosolic ROS production. Evidently, constant with other models of palmitate exposed cardiomyocytes [27-29], incubation of cells with doses ≥ 0.25 mM of this saturated FFA was enough to cause mitochondrial dysfunction and exacerbate oxidative stress-induced damage. However, it has also been clear that time of exposure with palmitate has been different across different investigations, including the current study, with some observing myocardial deterioration as early as 5 hours of exposure [27], and others reaching up to 24 hours [30, 31]. This may also suggest different concentrations of palmitate can be used to provoke different downstream effect associated with diverse disease complications.

Nevertheless, pertinent to the diabetic heart, lipid overload has been shown to severely deplete endogenous CoQ levels [13, 32], an element that has been rarely explored in experimental models of palmitate exposure. The current study showed that a dose-dependent increase in endogenous CoQ levels was up to a concentration of ≥ 5 mM of palmitate exposure, after which this component was reduced. In fact, palmitate doses > 5 mM caused severe mitochondrial toxicity, including reduction of cell viability and metabolic activity. Although endogenous CoQ levels were enhanced by palmitate doses ≤ 5 mM, this was not enough to counteract oxidative stress, but was sufficient to maintain cell viability of cardiomyocytes, suggesting a compensation mechanism. Interestingly, enhancement of CoQ levels with the lowest dose of palmitate (0.15 mM) was accompanied by a significantly reduction of CoQ oxidation status, as well as lower cytosolic ROS production. Thus, suggesting that additional studies are necessary to further investigate this aspect, especially establish the involvement of antioxidant defence mechanisms. The overall findings from this study strongly suggest that enhancement of endogenous CoQ levels in the myocardium may be an important adaptive mechanism to counteract oxidative stress, and maintain cell viability, as it was clear that as soon as CoQ levels were reduced at a concentration above 5 mM of palmitate, mitochondrial toxicity and cell viability were heavily impacted.

In summary, due to their abundance in the myocardium and their prominent role in oxidative metabolism, mitochondria have become ideal targets of intervention to protect against cardiovascular complications. In this case, critical understand of precise mechanisms implicated in enhanced myocardial lipid accumulation are necessary to protect against lipid-induced cardiotoxicity, such as that seen in DCM. From the overall findings, it appears that CoQ response may be crucial to improve mitochondrial function and thus protect against hyperlipidemia-induced insult. These results further suggest that therapeutic agents that can stimulate endogenous levels of CoQ may be beneficial in protecting the myocardium against diabetes associated.

## Abbreviations

ATP: adenosine triphosphate
CoQ9/10: Coenzyme Q9/10
DCFH-DA: 2’, 7’-dichlorfluorescein diacetate
DCM: diabetic cardiomyopathy
DMEM: Dulbecco’s Modified Eagle’s Medium
DPBS: Dulbecco’s Phosphate Buffered Saline
ECAR: extracellular acidification rates
FBS: fetal bovine serum
FFAs: free fatty acids
H_2_O_2_: hydrogen peroxide
HPLC: high-performance liquid chromatograph
O_2_^• −^: superoxide radical
•OH: hydroxyl radical
OCR: oxygen consumption rate
ROS: reactive oxygen species.

